# Probabilities of finding trace profile donors and their paternal relatives in Y-STR reference databases

**DOI:** 10.1101/2025.01.08.631054

**Authors:** Tóra Oluffa Stenberg Olsen, Niels Morling, Poul Svante Eriksen, Marie-Louise Kampmann, Helle Smidt Mogensen, Mikkel Meyer Andersen

## Abstract

Forensic investigative genetic genealogy using Y-chromosome short tandem repeat (Y-STR) DNA profiles can give investigative leads in criminal cases by searching for the Y-STR trace profile or similar but not identical Y-STR profiles in relevant Y-STR databases. We conducted a simulation study with Yfiler^™^ Plus and PowerPlex^®^ Y23 Y-STR profiles to estimate the probabilities of finding matches and near-matches in Y-STR databases. The success rate of finding the trace profile donors or their close relatives was quantified. We used the malan R software package to simulate the populations based on the Wright-Fisher model with the YHRD Y-STR mutation rates where uncertainties were incorporated in a Bayesian manner, a variance in reproductive success of 0.2, and a constant size for 100 generations followed by a 2% growth for 150 generations. Y-STR databases were generated by randomly drawing Y-STR profiles from a Yfiler^™^ Plus and PowerPlex^®^ Y23 Y-STR population data set, respectively.

In a population of 500,066 individuals, a database size of 0.5% of the population resulted in a Y-STR database match probability of ca. 6% and 10% for Yfiler^™^ Plus and PowerPlex^®^ Y23, respectively. Increasing the database size to 5% of the population resulted in a Y-STR match probability of ca. 41% and 54% for Yfiler^™^ Plus and PowerPlex^®^ Y23, respectively. When a Y-STR match was found in the database, the probability of one of the individuals with the matching profiles being related within five meioses to the trace donor was ca. 64% and 56% for Yfiler^™^ Plus and PowerPlex^®^ Y23, respectively, including the cases where the Y-STR profile originated from the donor. In this case, the closest relative in the database was found among the matching individuals with a probability of ca. 91%.

## 1. Introduction

Biological samples with DNA fragments collected for forensic investigations are primarily typed for autosomal short tandem repeat (STR) loci, and the STR profiles are compared to STR profiles of suspects or searched for matches in relevant databases [1]. If autosomal STR typing is unsuccessful and it is interesting for the case to distinguish between male and female donors, Y-STR typing may be relevant. The trace Y-STR profiles can be compared to those of suspects and searched for in relevant Y-STR databases.

Y-STR profiles are inherited unchanged from father to son, except for mutations at each locus [2, 3, 4, 5]. Commercial forensic Y-STR kits type for several Y-STRs, e.g., the Y-STR kit Yfiler^™^ Plus PCR Amplification Kit (Applied Biosystems) types 27 Y-STR loci [6]. The mutation probability in one or more Y-STR systems is several percent in all populations [2, 4]. Thus, genetic analyses based on results obtained with forensic Y-STR kits among male family members are simple but the analyses must include the mutation risk.

We explored the probability of finding (1) matching Y-STR profiles and Y-STR profiles similar but not identical to the trace profiles and (2) the trace donors of Y-STR matches and near-matching Y-STR profiles among the individuals who contributed to a Y-STR reference database and their paternal relatives in Y-STR databases.

## 2. Methods

The analyses were done in R [7] version 4.4.1 using the packages malan version 1.0.3 [8], tidyverse version 2.0.0 [9], and ggh4x version 0.2.8 [10]. The application code is available in [11].

### 2.1. Simulating populations using the malan R package

The procedure of simulating the populations using the malan R package was based on [12, 13]. The populations were constructed based on the Wright-Fisher model [14, 15], modified by the variance of the reproductive success of 0.2 to account for varying male reproductivity [12, 13]. We only considered the last three generations since only these individuals are assumed to be of interest in forensic cases. We refer to this population subset as the “live” population. We simulated “live” populations of 500,066 and 1,500,196 males 10 times with a constant size over 100 generations, followed by a 2% growth for each generation for 150 generations [12, 13].

We assigned Y-STR profiles of the Yfiler^™^ Plus kit [6] and the PowerPlex^®^ Y23 kit (Promega Corporation) [16], respectively, to the individuals in the simulated populations twice. DYS385a and DYS385b were considered independent loci [12, 13]. DYF387S1a and DYF387S1b were treated the same way. We only considered integer-valued alleles.

A founder Y-STR profile was assigned by randomly selecting a Y-STR profile from a reference database. For the Yfiler^™^ Plus kit, we used the profiles from Denmark in [17] and for the PowerPlex^®^ Y23 kit, we used the profiles from Denmark in [18]. Then the profile was transmitted to the founders’ male children. Potential one-repeat mutations with the allele mutation rates published at www.yhrd.org were introduced [2] where uncertainties were incorporated as in [12, 13]. We divided the DYS385 mutation rate equally between DYS385a and DYS385b [12, 13]. Simulated alleles outside the ranges of the allelic ladders were excluded. The process was repeated for each individual’s Y-STR profile in the following generations and continued until all individuals in the population were assigned a Y-STR profile.

### 2.2. “Cases” and reference databases

For each simulated “live” population, we drew 100 profiles corresponding to trace profiles. For each trace profile donor, 10 different reference databases of sizes 0.5%, 1%, 2%, and 5% of the “live” population were drawn randomly without replacement. Thus, for each population size and kit considered, 10 × 2 × 100 × 10 × 4 = 80,000 “cases” were created.

We calculated the number of one-repeat Y-STR mutations between the trace and the database profiles. For each “case”, the meiotic distance and the number of mutations between the trace profile and database profiles were registered. We registered the number of meioses up to 15 meioses and larger meiotic distances as more than 15 meioses.

## 3. Results

### 3.1. Probability of finding the trace profile and similar profiles in the database

We estimated the probability of observing the trace profile and profiles similar but not identical to the trace profile in a Y-STR reference database. For each “case”, we considered only one Y-STR profile with the smallest number of mutations to the trace profile.

Fig. 1 shows the probabilities of observing a given smallest number of mutations between the trace and database profiles in four databases of different fractions of two population sizes for each kit. The tendencies were similar for both population sizes. For the population size of 500,066, the database fractions 0.5%, 1%, 2%, and 5% of the population resulted in probabilities of the databases containing a profile matching the trace profile of ca. 6%, 12%, 22%, and 41%, respectively, for Yfiler^™^ Plus and ca. 10%, 19%, 32%, and 54%, respectively, for PowerPlex^®^ Y23. For the 0.5% database fraction, the highest probabilities were found at a minimum of four mutations. For the 2% database fraction for Yfiler^™^ Plus and the 1% database fraction for PowerPlex^®^ Y23, the probability of the smallest number of mutations was highest for zero mutations, i.e., a match, and decreased with increasing numbers of mutations.

**Figure 1:**
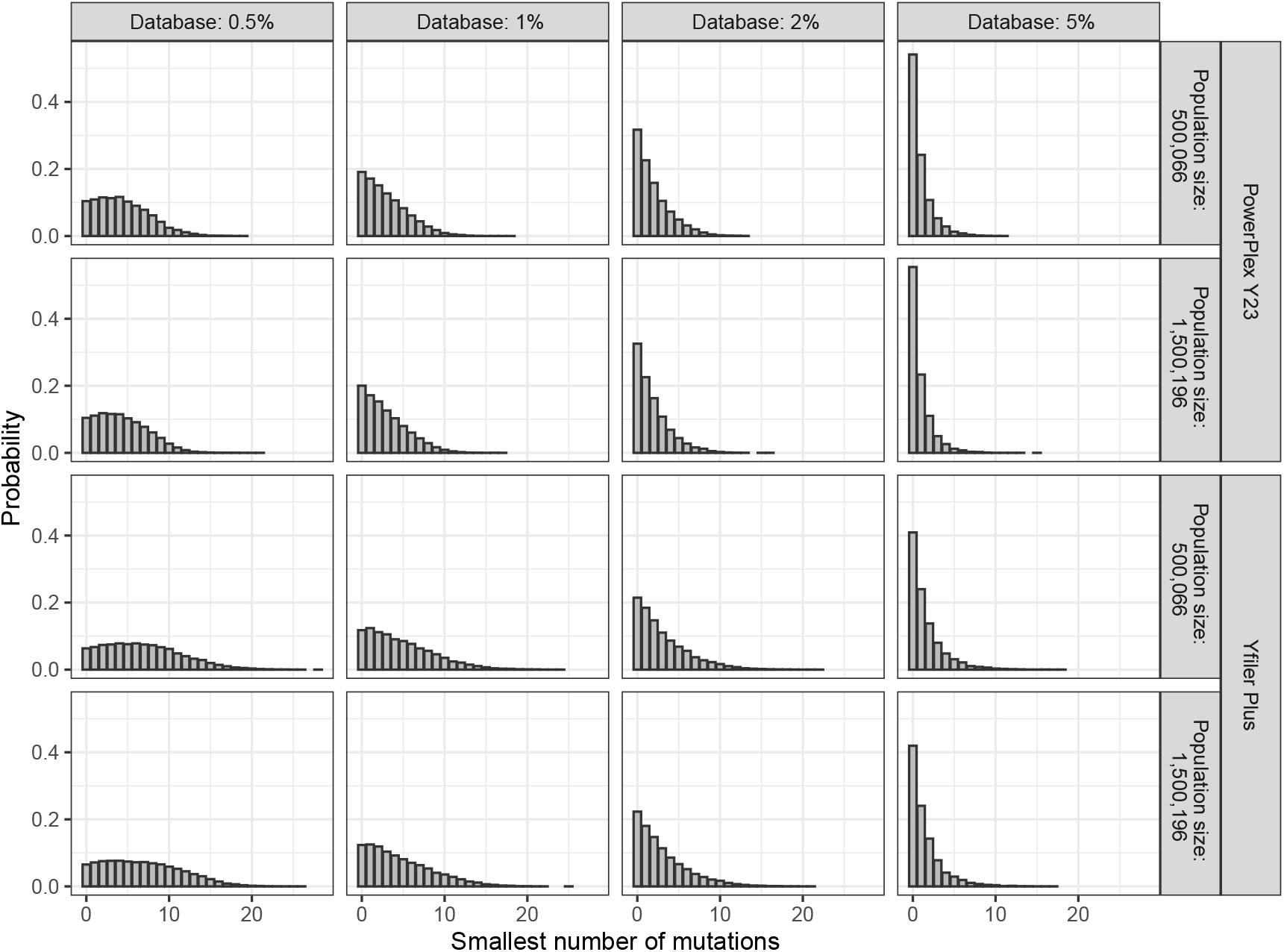
Probabilities of observing a given smallest number of mutations between the trace and database profiles in databases with 0.5%, 1%, 2%, and 5% of populations with 500,066 and 1,500,196 male individuals with PowerPlex^®^ Y23 and Yfiler^™^ Plus Y-STR profiles, respectively.

Fig. 2 shows the probabilities of observing matches and near-matches with one mutation between the trace and database profiles. For the 5% database fraction, the probability of finding a match was ca. 41% and 54% for Yfiler^™^ Plus and PowerPlex^®^ Y23, respectively, and ca. 24% for a near-match with one mutation for both kits.

**Figure 2:**
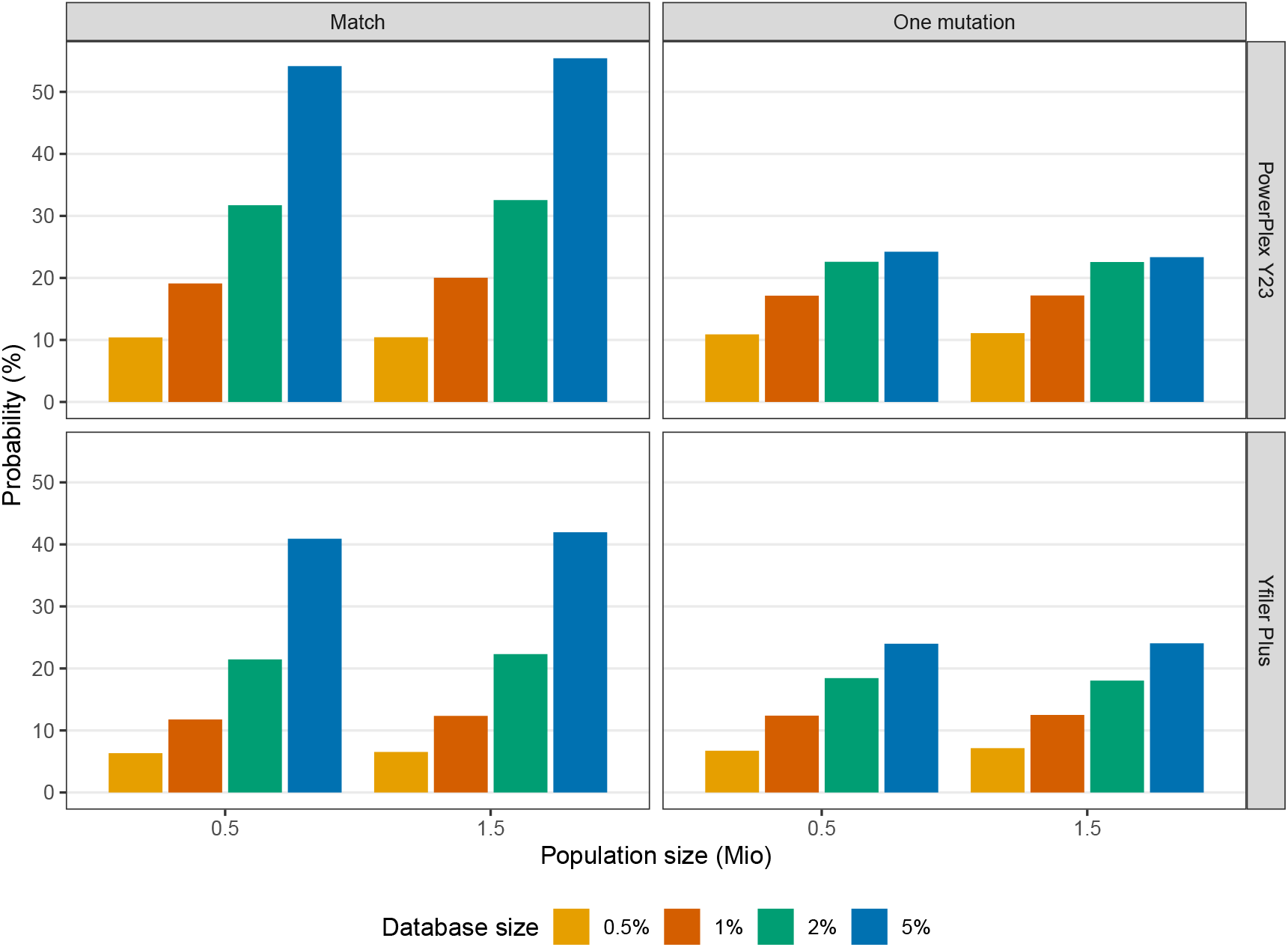
Probabilities of observing a match and one mutation near-match between the trace and database profiles in databases with 0.5%, 1%, 2%, and 5% of populations with 500,066 and 1,500,196 male individuals with PowerPlex^®^ Y23 and Yfiler^™^ Plus Y-STR profiles, respectively.

### 3.2. Probability of a match or a similar profile being from the donor or his paternal relative

Having examined the probability of observing the trace profile or profiles similar but not identical to the trace profile in the database, we analysed the relationship between the donors of the trace and database profiles. Among database profiles deviating by the same smallest number of mutations from the trace profile, we selected the profile with the smallest meiotic distance to the trace profile.

Fig. 3 shows the probabilities of observing Y-STR profiles from paternal relatives within meiotic distances of 5, 10, and 15, respectively, when the smallest number of mutations to the trace profile was zero, one, or two mutations between the trace profile and the trace donor’s or his paternal relatives’ profiles for each database fraction, population size, and kit. The probability of identifying the donor by match was ca. 7 − 12% with the four database fractions for Yfiler^™^ Plus and ca. 5 − 9% for PowerPlex^®^ Y23. The probability of identifying the donor of a trace or his close relatives within five meioses was ca. 48 − 63% for Yfiler^™^ Plus and ca. 34 − 56% for PowerPlex^®^ Y23 with the four database fractions, increasing to ca. 85 − 93% for Yfiler^™^ Plus and ca. 70 − 86% for PowerPlex^®^ Y23 for up to 15 meioses when the smallest number of mutations was zero, i.e., a match. The probability of observing a paternal relative’s profile among near-matching profiles with one mutation to the trace profile was between ca. 15% and 75% for Yfiler^™^ Plus and ca. 8% and 61% for PowerPlex^®^ Y23 depending on the meiotic distances and the database fractions. The corresponding probabilities for near-matching profiles with two mutations were from ca. 4% to 50% for Yfiler^™^ Plus and ca. from 1% to 32% for PowerPlex^®^ Y23.

**Figure 3:**
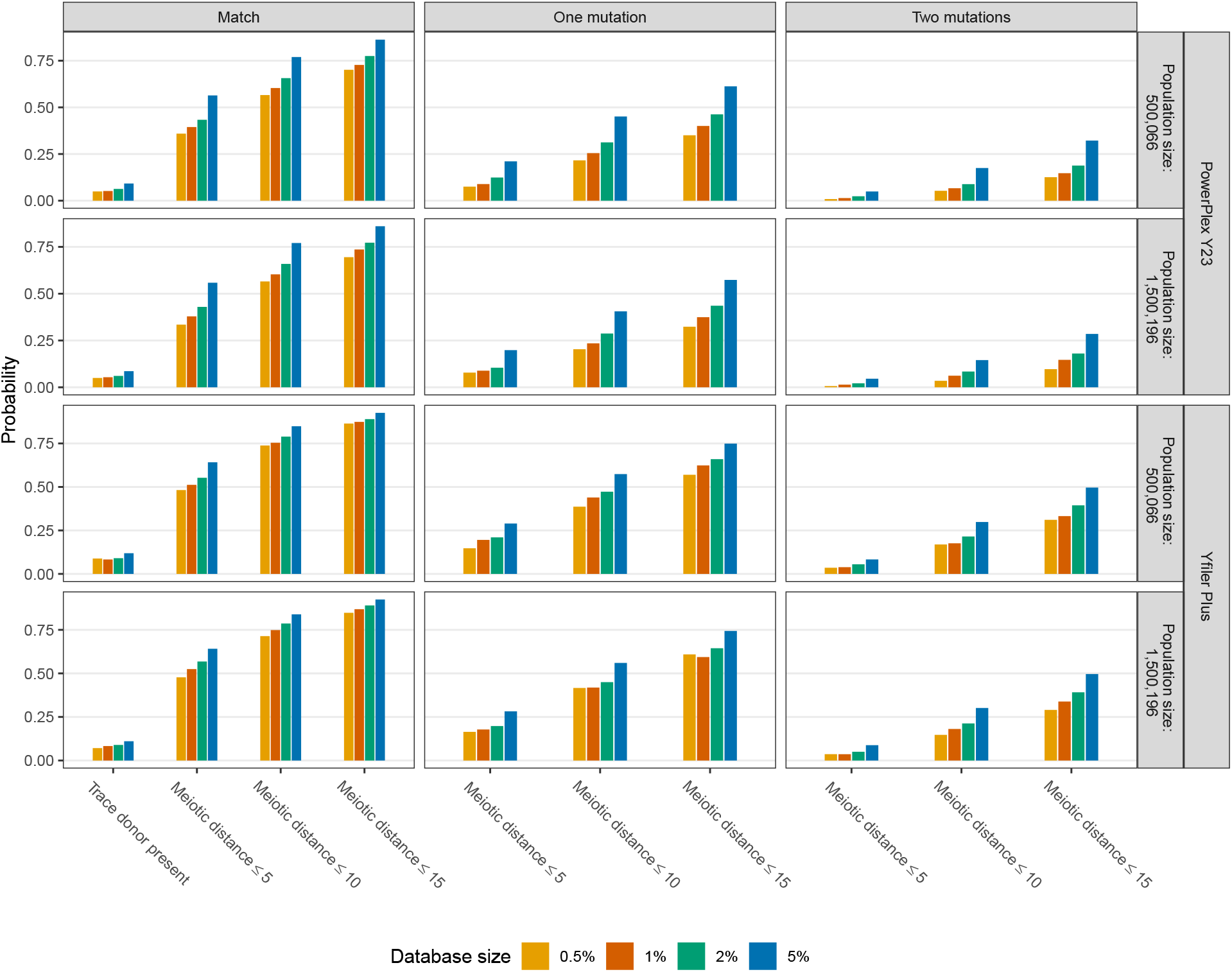
Probabilities of observing a Y-STR profile from a paternal relative within 5, 10, and 15 meioses to the trace profile when the smallest number of mutations between the trace and database profiles was zero, i.e., a match, one, or two mutations, respectively, in databases with 0.5%, 1%, 2%, and 5% of populations with 500,066 and 1,500,196 male individuals with PowerPlex^®^ Y23 and Yfiler^™^ Plus Y-STR profiles, respectively. If the smallest number of mutations was zero, the probability of a matching database profile being from the trace donor is also shown.

Increasing the size of the reference database increased the probability of an individual with a database profile with zero, one, or two mutations from the trace profile being a relative of the trace profile donor within a meiotic distance of 5, 10, and 15, respectively. This is most likely because increasing the size of the database increases the probability that the database contains multiple individuals with profiles with the smallest number of mutations to the trace profile, in which case we only consider one of the individuals having a profile with the smallest meiotic distance to the trace profile.

### 3.3. Probability of finding paternal relatives in the database

We considered the “cases” where the smallest number of mutations between the database and trace profiles was either zero or one, i.e., matches or one mutation near-matches because these are the most relevant ones.

Table 1 shows the probability of finding the closest paternal relative in the database. The majority of the closest relatives were found using this approach. For a database of 5% of a population of ca. 0.5 million male individuals where one profile in the database matched the trace profile, the probability of finding the closest paternal relative to the trace donor in the database was ca. 91% for both kits. Increasing the database size slightly decreased the probability of finding the trace donor’s closest relative in the database.

**Table 1:** Probabilities of finding the profile of the closest paternal relative to the trace profile donor among the database profiles and probabilities of finding more than one and three Y-STR profiles, respectively, with the smallest number of mutations of one or zero, i.e., a match, and with at least one such profile in the database and one profile in the reference database within a meiotic distance up to 15 to the trace profile. Notice that 0.00 is used for positive values less than 0.005, and 0 is used when no “cases” occurs.

|  | Population size | Database size (%) | Probability of finding the closest relative in the database |  | Probability of observing more than one database profile |  | Probability of observing more than three database profiles |  |
| --- | --- | --- | --- | --- | --- | --- | --- | --- |
|  |  |  | Match | One mutation | Match | One mutation | Match | One mutation |
| PowerPlex® Y23 | 500,066 | 0.5% | 0.98 | 0.96 | 0.10 | 0.13 | 0.00 | 0 |
|  | 500,066 | 1% | 0.96 | 0.95 | 0.18 | 0.21 | 0.01 | 0.01 |
|  | 500,066 | 2% | 0.93 | 0.90 | 0.32 | 0.34 | 0.03 | 0.04 |
|  | 500,066 | 5% | 0.91 | 0.88 | 0.53 | 0.54 | 0.16 | 0.18 |
|  | 1,500,196 | 0.5% | 0.98 | 0.97 | 0.11 | 0.11 | 0.00 | 0 |
|  | 1,500,196 | 1% | 0.96 | 0.96 | 0.20 | 0.22 | 0.00 | 0.01 |
|  | 1,500,196 | 2% | 0.93 | 0.91 | 0.33 | 0.35 | 0.04 | 0.04 |
|  | 1,500,196 | 5% | 0.91 | 0.87 | 0.55 | 0.55 | 0.16 | 0.18 |
| Yfiler™ Plus | 500,066 | 0.5% | 0.97 | 0.97 | 0.07 | 0.08 | 0 | 0 |
|  | 500,066 | 1% | 0.96 | 0.95 | 0.12 | 0.13 | 0.00 | 0.00 |
|  | 500,066 | 2% | 0.95 | 0.92 | 0.22 | 0.22 | 0.01 | 0.02 |
|  | 500,066 | 5% | 0.91 | 0.88 | 0.39 | 0.42 | 0.06 | 0.08 |
|  | 1,500,196 | 0.5% | 0.98 | 0.96 | 0.06 | 0.08 | 0.00 | 0.00 |
|  | 1,500,196 | 1% | 0.96 | 0.94 | 0.11 | 0.13 | 0.00 | 0.00 |
|  | 1,500,196 | 2% | 0.94 | 0.91 | 0.21 | 0.26 | 0.01 | 0.01 |
|  | 1,500,196 | 5% | 0.91 | 0.87 | 0.40 | 0.43 | 0.06 | 0.08 |

Table 1 shows the probability of observing more than one and three database individuals, respectively, whose profiles matched the trace profiles or deviated by one mutation. Generally, a maximum of three individuals’ profiles in the databases matched or deviated by one mutation from the trace profile for Yfiler^™^ Plus Y-STR profiles. For PowerPlex^®^ Y23 Y-STR profiles and a 5% database fraction, more than three matches occurred in ca. 16% of the databases containing at least one matching profile and more than three one mutation near-matches occurred in ca. 18% of the databases containing a profile deviating by one mutation.

## 4. Discussion

The success of obtaining investigative leads in criminal cases by family searching in Y-STR databases depends on the probability of finding matches or near-matches between the trace’s and database individuals’ Y-STR profiles. The degree of relationship between the database and trace donor is quantified by the profiles’ meiotic distance, estimated by the mutation number.

We created a simulation model based on Yfiler^™^ Plus and PowerPlex^®^ Y23 Y-STR allele and mutation frequencies observed in European populations collected by the YHRD [2]. We used a simple mutation model, considering only integer-valued alleles and assuming mutations of only one STR-repeat unit (one-step mutations), although mutations of two or more steps are observed with a much smaller frequency [19]. Possible back-mutations were not taken into consideration. We simulated populations of ca. 0.5 and 1.5 million male individuals and four databases representing 0.5%, 1%, 2%, and 5% of each population size. We estimated the probabilities of finding matches and near-matches with up to 15 meioses between the trace and database profiles for all databases.

The population size did not influence the success rate in the databases much. For a population size of 500,066 and a database of 5% of the population, the probability of the database containing a profile matching the trace profile was ca. 41% and 54% for Yfiler^™^ Plus and PowerPlex^®^ Y23, respectively (Fig. 1 and 2). If a match between the database and trace profiles was found, the probability of finding the trace donor within five meioses of the database individual was ca. 64% and 56% for Yfiler^™^ Plus and PowerPlex^®^ Y23, respectively (Fig. 3). Thus, the probability of the database containing a matching profile from a male related to the trace donor within five meioses was ca. 26% and 30% for Yfiler^™^ Plus and PowerPlex^®^ Y23, respectively. If at least one database profile matched the trace profile, the probability of finding the trace donor’s closest paternal relative in the database among the matches in the database was ca. 91% for both kits (Table 1). When at least one match was found, the probability of observing more than one match was ca. 39% and 53% for Yfiler^™^ Plus and PowerPlex^®^ Y23, respectively, and the probability of observing more than three matches was ca. 6% and 16% for Yfiler^™^ Plus and PowerPlex^®^ Y23, respectively.

The probability of the smallest number of mutations between the trace and database profiles being one was ca. 24% for both kits (Fig. 1 and 2). When a one mutation near-match between the database and trace profiles was found, the probability of finding the trace donor within five meioses of the database individual was ca. 29% and 21% for Yfiler^™^ Plus and PowerPlex^®^ Y23, respectively (Fig. 3). Thus, the probability of observing a one mutation near-match with the profile of a paternal relative of the trace donor within five meioses was ca. 7% and 5% for Yfiler^™^ Plus and PowerPlex^®^ Y23, respectively.

The probability of finding paternal relatives’ profiles in Y-STR databases is increased if the size of the reference database is increased. The probability of finding profiles of distantly related individuals is also increased. If the Y-chromosome analysis is to be used to analyse male relationship beyond approximately five meioses, the information in the genetic analyses must be increased, e.g., by using more Y-STR or Y-SNP markers.

